# *LatenZy*, non-parametric, binning-free estimation of latencies from neural spiking data

**DOI:** 10.1101/2025.06.30.662308

**Authors:** Robin Haak, J. Alexander Heimel

**Author notes:** **Contact information** Correspondence should be addressed to J.A.H.

## Abstract

Precisely estimating the onset of neural spiking responses and the timing at which activity begins to diverge between conditions is crucial for understanding temporal dynamics in brain information processing. Conventional methods require arbitrary parameter choices such as bin widths and response thresholds, limiting reproducibility and comparability. Here, we present *latenZy* and *latenZy2*, two non-parametric, binning-free methods that directly analyze spike times using cumulative statistics and iterative refinement, without assumptions about response shape. *LatenZy* estimates neuronal response onset latency, while *latenZy2* detects when spiking activity diverges between conditions. We validate these methods on electrophysiological datasets from mouse and macaque visual cortex, and show that they outperform standard approaches in precision, robustness, sensitivity, and statistical power. *LatenZy* captures contrast-dependent latency shifts and hierarchical timing across visual areas, and *latenZy2* reveals earlier attentional modulation in higher visual cortex consistent with top-down feedback. Together, they offer scalable, parameter-free tools for reliable latency estimation in large-scale neural recordings. Open-source implementations are available in Python and MATLAB.

## Introduction

Precise quantification of *when* spiking activity is modulated by internal or external events is essential in neurophysiological studies. A widely used metric is response latency – the time between an event and the onset of a neural response (1,2). Response latencies reveal the temporal structure of neural activation, allowing researchers to infer causal relationships between stimuli, neural circuits, and behavior (3–5). By identifying the moment a neuron or brain region becomes active, one can distinguish potential drivers of a response from those involved in later, downstream processing. Moreover, comparing latencies across neurons and areas provides critical insights into the functional hierarchy and information flow within the brain (6–9). In many experimental paradigms, it is equally important to determine when spiking activity begins to diverge between conditions. This latency marks the onset of selective neural coding, which is critical for understanding processes like sensory discrimination (10–12). Consequently, latency estimation techniques have been a central focus of technological development over the past decades.

Binning spike trains—creating a peri-event time histogram (PETH)— is the first step in many existing latency estimation methods (1,2,13–17), which requires choosing an appropriate bin width. Large bins obscure fine temporal features, while small bins may amplify noise, both of which can reduce the accuracy and reproducibility of latency estimates. The optimal bin width depends on factors such as a neuron’s firing rate and the smoothness of its event-related modulation (18–20)—properties that vary widely across neurons—and is influenced by experimental factors, such as the number of stimulus repetitions. Additionally, most methods rely on choosing arbitrary parameters, such as response thresholds or minimum response durations. These choices complicate meaningful comparisons of latency estimates across neurons, brain regions, and studies. With large-scale electrophysiological recordings now commonplace (21–25), there is a growing need for methods that circumvent these arbitrary parameter choices.

To address key limitations in existing latency estimation approaches, we developed *latenZy* and *latenZy2*, two novel, binning-free methods for estimating the onset of neural spiking activity. *LatenZy* builds upon a previously developed framework for detecting event-locked changes in spiking activity (26,27), adapting it to estimate when neural responses begin following discrete events with high temporal precision. *LatenZy2* extends this approach to determine when spiking activity begins to diverge between experimental conditions. Both methods leverage cumulative spike timing and iterative refinement to ensure robust and accurate latency estimates across diverse response profiles. We show that *latenZy* and *latenZy2* outperform traditional PETH-based approaches, recovering known physiological effects such as contrast-dependent latency shifts and hierarchical timing differences, while maintaining greater statistical power at smaller sample sizes. Together, these tools provide a unified, binning-free framework for precise and reliable estimation of neural response latencies in both stimulus-evoked and condition-specific contexts.

## Results

### *LatenZy*: binning-free estimation of neural response latencies

Neurophysiological studies often require estimation of neural response latency—the moment neural activity begins to change relative to an event (1,2). Many standard latency estimation methods rely on arbitrary parameter choices, like the selection of a bin width or statistical threshold, that can compromise both accuracy and reproducibility. To overcome this, we developed *latenZy*, a method that avoids binning and other parameter choices altogether. *LatenZy* is an extension of the Zenith of Event-based Time-locked Anomalies (ZETA) test (26,27) and follows many of the same core steps. It is designed to estimate when a neural response begins, defined as the first significant, time-locked change in spiking activity within a time window around an event, such as a stimulus onset. This time point is identified through an iterative procedure that progressively refines the estimate (Fig. 1A). A detailed description of *latenZy* is provided in the Methods section; here, we offer a brief overview of its procedure.

**Figure 1.**
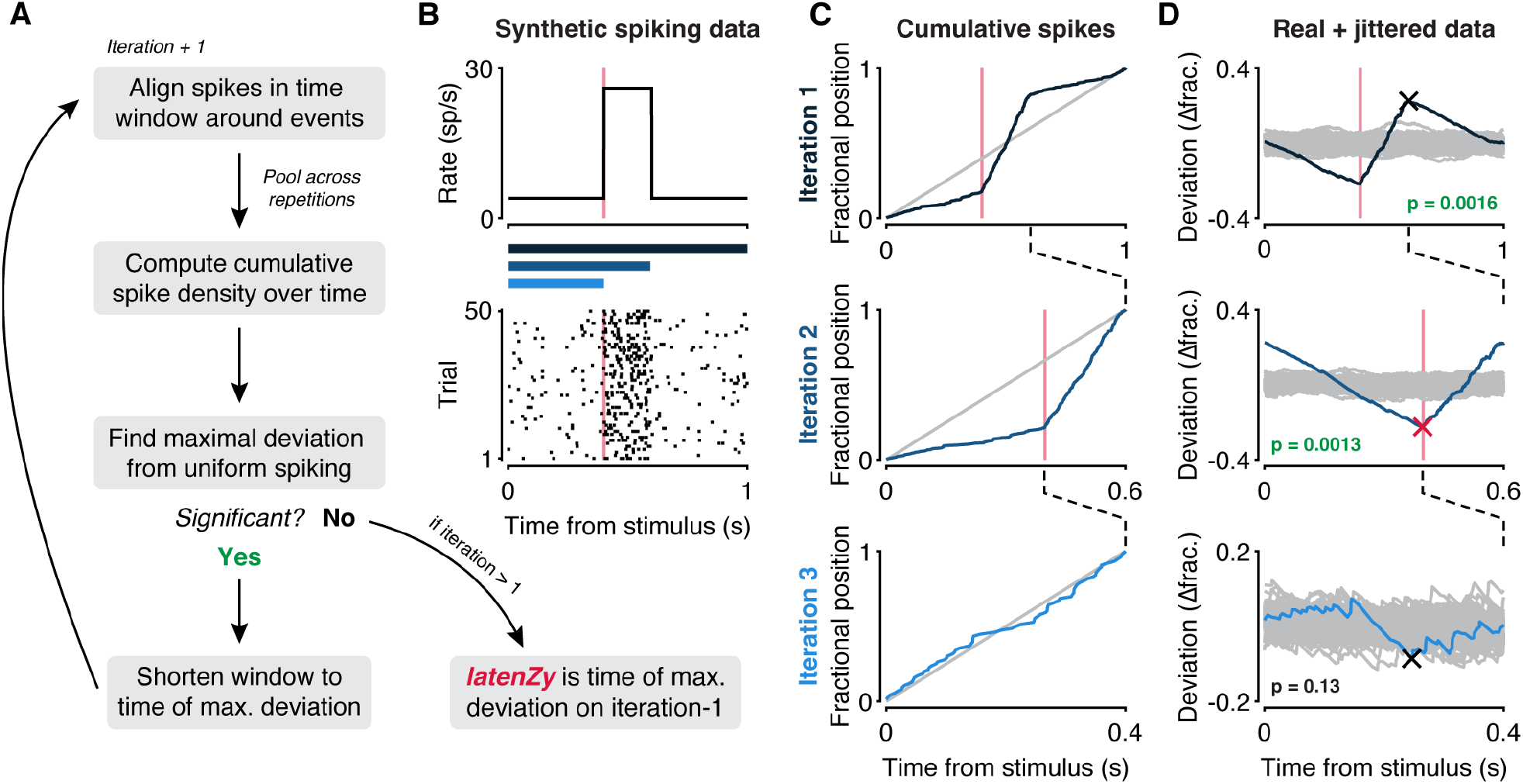
Overview of the *latenZy* method. (A) Block diagram illustrating one iteration of *latenZy*. (B) To illustrate its procedure, *latenZy* was applied to spiking activity generated using an inhomogeneous Poisson process. The top panel shows the time-varying response function used to drive the simulations, while the bottom panel is a raster plot of the resulting spike trains aligned to stimulus onset. The vertical pink lines indicate the ground-truth response onset, and the blue bars represent the time windows evaluated in each iteration of *latenZy*, corresponding to the rows in C and D. (C) *LatenZy* avoids binning by using the spike times to construct a fractional position for each spike (blue curve) and compares this with a linear baseline (gray line). (D) In each iteration, the maximal deviation from the linear baseline (blue curve) is determined (cross) and its significance is determined by comparing it to the variability over repeats of the procedure with jittered event times (gray curves). If significant (p < 0.05), the upper bound of the time window is shortened to the time point corresponding to the maximal deviation, and the steps are repeated. This continues until no further time-locked spiking is detected—indicating that the response latency was determined on the previous iteration. The red cross indicates the *latenZy* estimate, which closely matches the ground-truth response onset in this example.

We start by aligning spikes within a window surrounding each event time, like constructing a raster plot (Fig. 1B, bottom). By pooling spikes across the event repetitions, we generate a single vector of spike times relative to the events and compute their cumulative distribution over time (Fig. 1C, blue curves). We then identify the point of maximal deviation from a linear baseline and its associated latency (Fig. 1D, crosses). Next, we assess the significance of the time-locking through multiple bootstraps by jittering event times to create a null-hypothesis distribution (Fig. 1D, gray curves). The maximal deviation is scaled relative to this null distribution and transformed into a z-score and associated p-value using a Gumbel distribution approximation (26). If this deviation is statistically significant (p < 0.05, default cutoff), the upper limit of the time window used for analysis is shortened to the estimated latency, and the procedure is repeated iteratively. This process continues until no further significant modulation is detected. The *latenZy* estimate is the earliest time point from the start of the window where significant, time-locked modulation of spiking activity is detected through this iterative process (Fig. 1D, red cross).

The theoretical foundation of *latenZy* relies on the behavior of the cumulative distribution of spike times pooled across repeated trials. In the limit of many trials, variability in spike timing averages out, and during periods of purely spontaneous activity, the cumulative distribution converges to a linear function. This linearity means that any maximal deviation must occur at the boundaries of the spontaneous interval. Since the lower bound of the analysis window is fixed and anchored at zero deviation, the maximal deviation within the spontaneous segment must occur at its upper bound—marking the transition from spontaneous to evoked, time-locked spiking. *LatenZy* exploits this by iteratively trimming the analysis window from its upper bound, progressively excluding time points already identified as significantly modulated. The iteration immediately preceding the final one thus contains the critical transition where the slope of the cumulative distribution first begins to change. By detecting the last statistically significant deviation associated with this boundary, *latenZy* converges on a principled and accurate estimate of neural response latency.

Although this provides the theoretical basis for *latenZy*, its true strength lies in how well it performs on real neural data, as we demonstrate below. Instead of relying on idealized assumptions—like neurons spiking with Poisson statistics—*latenZy* uses the ZETA test’s shuffle-based approach to assess significance directly from the data (26,27). By comparing observed spike timing to a null distribution created by jittering event times, *latenZy* accounts for the actual variability in real datasets. This makes it especially well-suited for real, non-Poissonian neural activity.

To illustrate the effectiveness of *latenZy*’s binning-free approach for accurate and robust latency estimation across varied neural response patterns, we applied it to spike trains recorded from neurons in the zebrafish retina over multiple trials of light flash stimulation. These examples include a neuron with a relatively high spiking rate and a strong transient response (Fig. 2A), a neuron with a low rate and a brief response (Fig. 2B), and one high-rate neuron exhibiting complex dynamics characterized by initial suppression followed by multiple response peaks (Fig. 2C). The absence of binning in *latenZy* enables preservation of high temporal precision, resulting in latency estimates that correspond closely with visually identifiable response onsets in the raster plots (Fig. 2A-C, left, red vertical lines). In the next sections, we evaluate *latenZy*’s performance against two established bin-based latency estimation methods.

**Figure 2.**
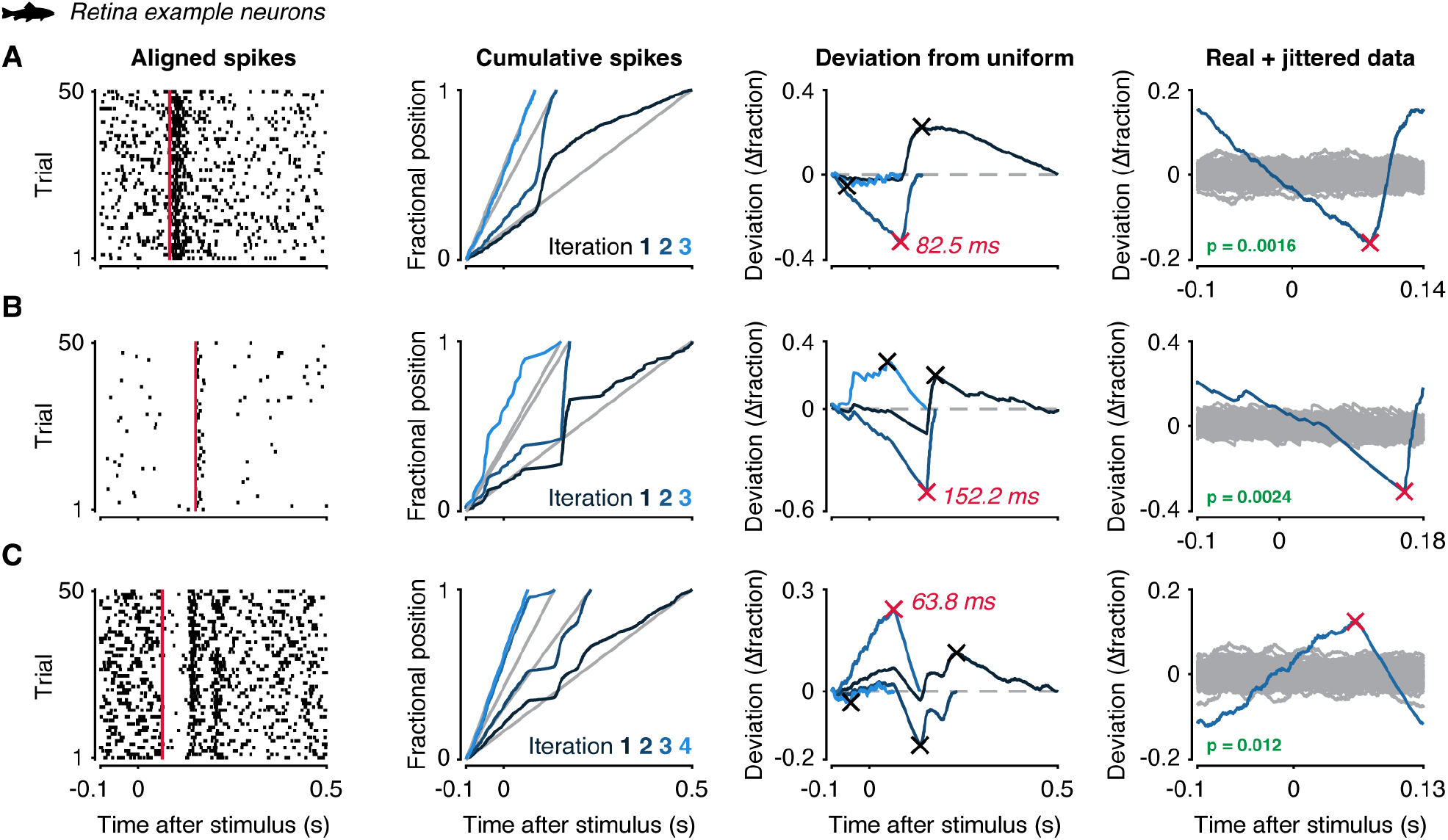
Application of *latenZy* to example neurons recorded from zebrafish retina. *LatenZy* was applied to spike trains from three individual neurons that exhibited distinct spiking patterns in response to a light flash: (A) a high-rate neuron (31.9 sp/s) exhibiting a strong transient response, (B) a low-rate neuron (3.5 sp/s) with a brief response, and (C) a high-rate neuron (52.1 sp/s) showing complex dynamics. For each neuron, the plot (from left to right) show: the spike trains aligned to light flash onset; the fractional spike position curves (blue) with linear baselines (gray) computed across iterations; the deviations from a uniform firing rate, with maximal deviations indicated by crosses; and the mean-subtracted deviation curve (blue) from the iteration in which the *latenZy* estimate was obtained (red cross), shown alongside the jittered data (gray curves) used to assess statistical significance in the penultimate iteration. For each neuron, the *latenZy* estimate (red vertical line) closely matches the visually apparent response onset in the raster plots (left).

### *latenZy* provides more stable and robust latency estimates than bin-based methods

We applied *latenZy* to 686 mouse primary visual cortex (V1) neurons responding to drifting gratings (Fig. 3A, B, example neuron), obtaining a median latency estimate of 61.7 ms (Fig. 3F, inset, red line)—well within the range reported in previous studies (9,28–31). To benchmark *latenZy*, we compared it to two commonly used, PETH-based latency estimation approaches: the half-max (15,16) and standard deviation (SD)-threshold methods (13) (Fig. v3C). The half-max method defines latency as the center of the first bin in which the firing rate deviates from baseline by at least half the amplitude of the peak response, in either direction (baseline ± 0.5 × |peak|). The SD-threshold method defines latency as the center of the first bin in which the firing rate exceeds the baseline by *k* standard deviations (baseline ± *k* × baseline SD). We use *k* = 2.58, a common choice corresponding to a two-tailed significance level of 0.01 (32,33), but other arbitrary choices for *k* are also frequently used. We selected these methods for their widespread use and for reflecting complementary principles: half-max captures large, relative changes in response magnitude, while SD-threshold identifies statistically significant deviations from baseline variability. Moreover, by collapsing spike times over trials and ignoring trial variability, these PETH-based methods share a key assumption with *latenZy*, supporting their use as benchmarks.

**Figure 3.**
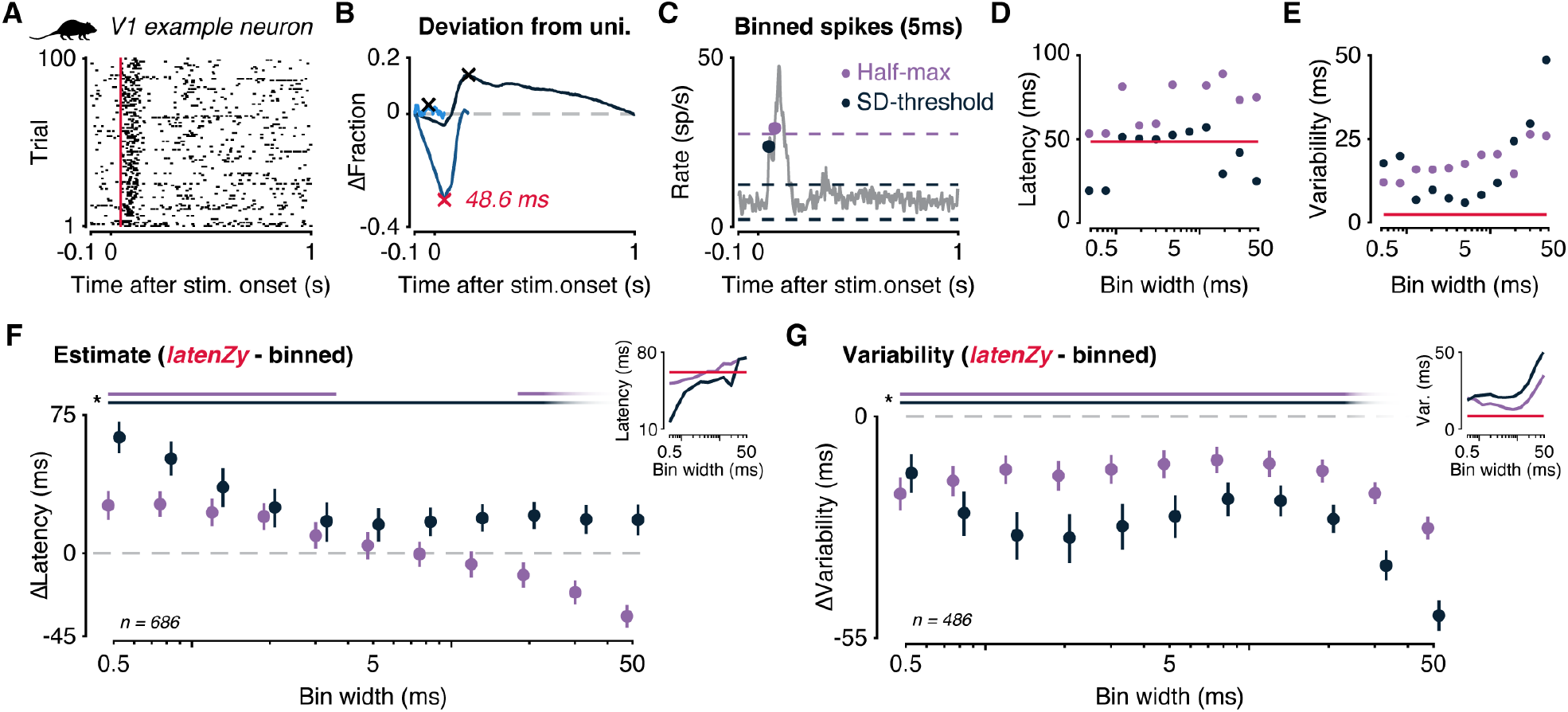
*LatenZy* provides more stable and precise latency estimates than bin-based methods. (A) Raster plot of an example mouse V1 neuron aligned to the onset of a drifting grating stimulus. Red vertical line indicated the *latenZy* estimate. (B) Corresponding *latenZy* analysis illustrating deviation from a uniform firing rate; the estimated latency is marked with a red cross. (C) *LatenZy* was compared against two conventional bin-based latency estimation methods: the half-max method, which identifies latency as the first bin reaching half the peak response, and the SD-threshold method, which defines latency as the first bin exceeding baseline firing by 2.58 standard deviations. (D) All methods were applied across a range of bin widths (0.5–50 ms) to quantify latency estimates, and resampling-based variability was assessed (E). (F) Bootstrapped comparisons revealed that *latenZy*’s estimates generally diverged from both bin-based methods, except at intermediate bin widths (5–12.5 ms) for the half-max method, where convergence was observed. Bin-based estimates varied substantially with bin width. Inset shows the median latency estimates for each method and bin width. (G) While absolute latency depends on each method’s definition, comparing estimate variability provides insight into reliability. *latenZy* consistently showed lower variability than both bin-based methods across all bin widths tested. Error bars represent mean with 95% CI. Lines at the top indicate significance (*, p < 0.05).

We applied the half-max and SD-threshold methods across a range of bin widths (0.5 to 50 ms), obtaining latency estimates that clearly depend on the bin width used (Fig. 3D). For short and long time bins, the SD-threshold method produced unrealistically fast onsets. We also quantified the variability of the latency estimates by repeated subsampling of the trials (Fig. 3E). We repeated this analysis for all V1 neurons and used bootstrapping to generate distributions of the mean differences between *latenZy* and each bin-based method (*latenZy* minus bin-based) for statistical comparison. The latency estimates of both bin-based methods varied considerably as a function of bin width, particularly for the SD-threshold method, which produced artefactually early latencies at small bin widths (Fig. 3F, inset). *LatenZy*’s estimates were different from those produced by both bin-based methods (p < 0.05), with the exception of the half-max method at intermediate bin widths (5 to 12.5 ms), where estimates are similar (Fig. 3F).

Although absolute latency values vary according to each method’s definition of response onset, examining the variability of latency estimates offers valuable complementary insight into their reliability and precision. In this context, *latenZy* consistently demonstrated lower variability than both bin-based methods across all tested bin widths (p < 0.05; Fig. 3G), highlighting its robustness and reproducibility in detecting response onset timing.

### *LatenZy* outperforms alternative methods in capturing contrast-driven latency shifts

Because there is no direct ground truth for neuronal response latencies, benchmarking latency estimation methods requires leveraging well-established physiological relationships. One such relationship is modulation of visual cortical response latencies by stimulus contrast, where higher contrast typically elicits faster responses (15,16,34,35). To evaluate whether *latenZy* captures this relationship more accurately than existing bin-based methods, we applied it alongside the half-max and SD-threshold methods to a publicly available dataset of spiking responses from mouse V1 neurons stimulated with drifting gratings at multiple contrast levels (9) (Fig. 4A, B).

**Figure 4.**
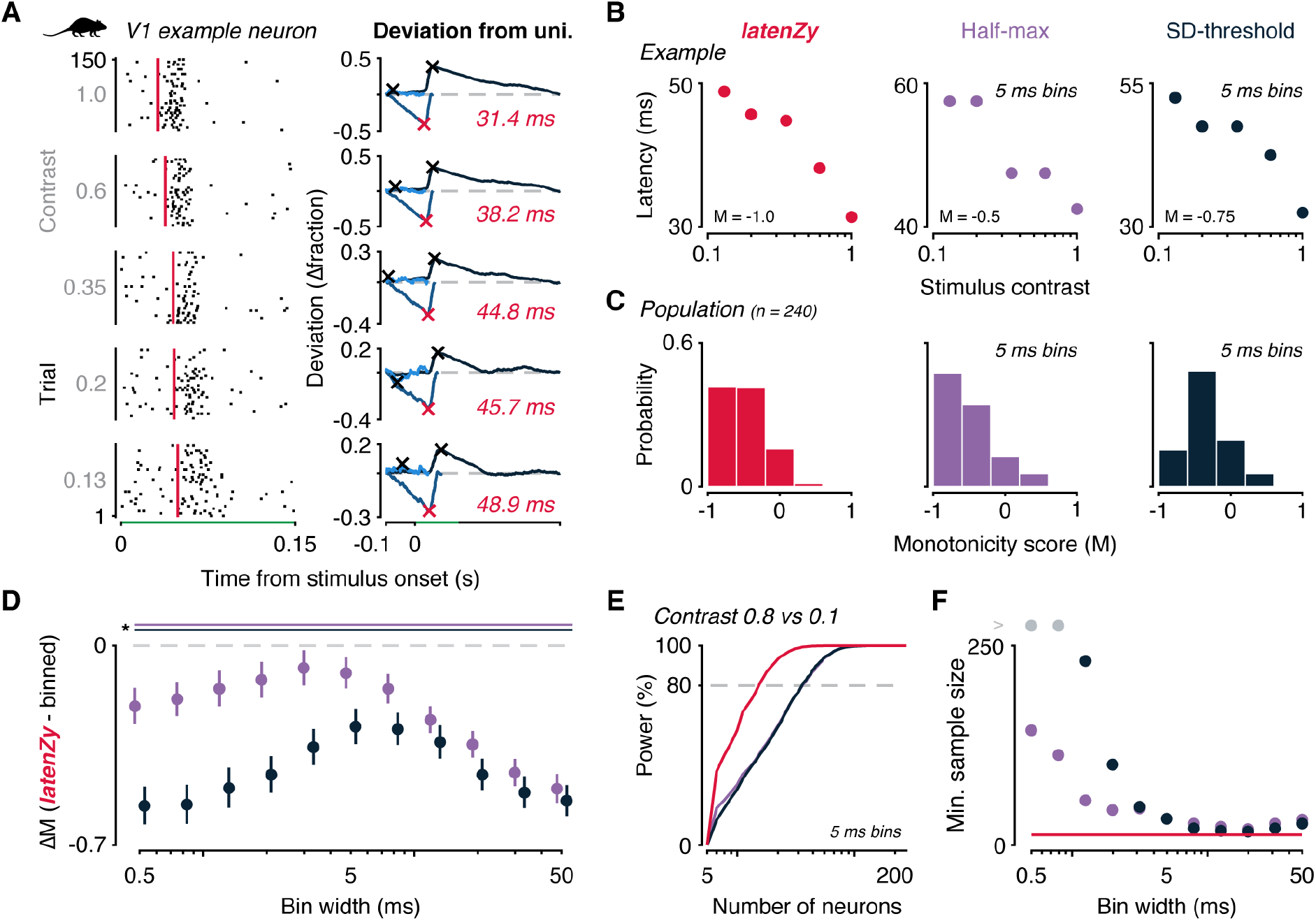
*LatenZy* outperforms bin-based methods in capturing contrast-dependent latency shifts. (A–B) *LatenZy* was benchmarked against bin-based methods using the known relationship between stimulus contrast and neural response latency, where higher contrast typically elicits earlier responses. Spike trains from a mouse V1 neuron exposed to drifting gratings at multiple contrast levels were analyzed using *latenZy*, half-maximum, and SD-threshold methods. Red vertical lines indicate the *latenZy* estimates. A monotonicity score (MS), where –1 indicates perfect monotonicity, quantified the consistency of latency decreases across contrasts. For an example neuron (B, same as A), *latenZy* produced a strictly monotonic trend, whereas bin-based methods showed ties due to bin overlap. (C) At the population level, both *latenZy* and the half-max method (5 ms bins) effectively captured the expected contrast-latency relationship, whereas the SD-threshold method showed reduced sensitivity. (D) Bootstrapped comparisons confirmed *latenZy*’s consistently higher monotonicity scores across all tested bin widths. Error bars represent mean with 95% CI. Lines at the top indicate significance (*, p < 0.05). (E–F) Power analyses using Wilcoxon signed-rank tests (high contrast: 0.8; low contrast: 0.1) showed that *latenZy* required fewer neurons to reach 80% power at α = 0.05, both at 5 ms bins (E) and across smaller bin widths (F).

We quantified the relationship between latency and contrast for each neuron using a strict monotonicity score, where −1 indicates a strictly monotonic decrease in latency with increasing contrast, and 1 indicates a strictly monotonic increase. For an example neuron, *latenZy* yielded a perfectly monotonic decrease, whereas the binned methods produced ties— cases where two contrast levels yielded indistinguishable latency estimates due to overlapping bins (Fig. 4B). At the population level, both *latenZy* and the half-max method, using a 5 ms bin width, effectively captured the expected monotonic decrease, while the SD-threshold method showed weaker alignment at the same bin width (Fig. 4C).

To assess how bin width affects the relative performance of latenZy and bin-based methods, we compared their monotonicity scores across a range of bin widths Because *latenZy* operates without binning, its estimates remain constant, while those from bin-based methods vary with bin width. Using bootstrapping, we generated distributions of mean differences in monotonicity scores between *latenZy* and each bin-based method (*latenZy* minus bin-based). Across all tested bin widths, *latenZy* consistently outperformed both bin-based approaches, yielding significantly more negative monotonicity scores (p < 0.05; Fig. 4D). These findings demonstrate that *latenZy* offers more reliable and sensitive detection of contrast-dependent latency shifts, with robustness unaffected by binning choices.

To compare latency estimates across methods, we performed a power analysis using Wilcoxon signed-rank tests to detect differences in response latencies between two contrast levels (0.8 vs. 0.1). With *latenZy* estimates, we achieved 80% power at a significance level of 0.05 with fewer neurons than with tests based on either bin-based method at a 5 ms bin width (Fig. 4E) and across smaller bin widths (Fig. 4F), indicating greater sensitivity to latency differences even with smaller sample sizes. Although this advantage diminished at larger bin widths, *latenZy* consistently required slightly fewer neurons to reach statistical significance. These results suggest that *latenZy* provides more sensitive and reliable latency estimates for detecting contrast-dependent shifts than bin-based alternatives.

### *LatenZy* captures the hierarchy of mouse visual cortex better than bin-based methods

The mammalian visual system is organized in a hierarchical manner, with information progressing through a series of interconnected cortical areas (36). A previous study (37) assigned anatomical hierarchy scores to individual areas in the mouse visual cortex by analyzing inter-area connectivity to determine the optimal network structure. In their analysis, V1 received the lowest hierarchy score, the rostrolateral area (RL), lateromedial area (LM), and anterolateral area (AL) were assigned intermediate scores, and the posteromedial area (PM) and anteromedial area (AM) ranked highest among the regions considered (Fig. 5A, B). Subsequent studies have validated the functional significance of these anatomical hierarchy scores by showing strong correlations with stimulus-evoked response latencies and other dynamical measures (9,29,38). Here, we use these anatomical hierarchy scores as a reference framework for evaluating the performance of *latenZy*.

**Figure 5.**
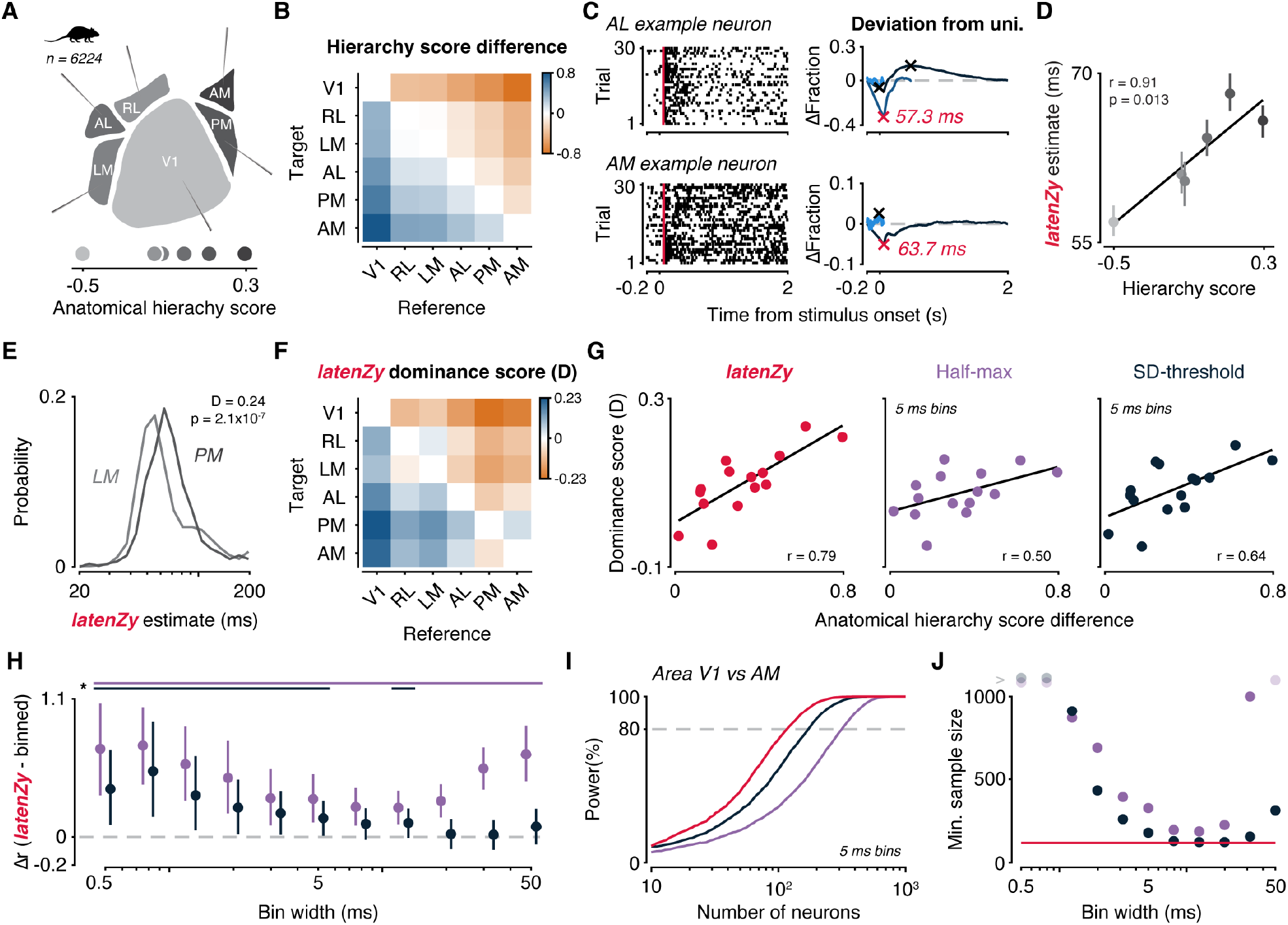
*LatenZy* captures meaningful latency differences that align with anatomical hierarchy across mouse visual cortex. To assess whether *latenZy* captures meaningful timing differences, we leveraged the known anatomical hierarchy of mouse visual cortex. (A) Hierarchy scores derived from anatomical tracing data (37) are shown for six cortical areas. (B) Replotted as pairwise differences, these scores provide a reference for expected temporal ordering between areas. (C) *LatenZy* was applied to neurons recorded from different regions, including two example cells shown here. (D) Median *latenZy* estimates across areas showed a strong correlation with anatomical hierarchy scores, demonstrating that *latenZy* accurately reflects the expected progression of visual processing stages. Error bars indicate bootstrapped 95% CI. (E) Distributions of *latenZy* estimates for two areas illustrate the shift in response timing across the hierarchy. (F) To more directly quantify relative timing, we computed latency dominance scores between all pairs of areas, capturing how often neurons from one area responded earlier than another. For example, a score of 0.24 for the PM vs. LM pair (E) indicates that PM neurons tend to have longer response latencies than LM neurons. (G) These scores showed strong correspondence with the anatomical hierarchy differences, outperforming bin-based estimates at a 5 ms resolution. (H) Bootstrapped comparisons across bin widths confirmed that *latenZy* consistently tracked the anatomical structure more closely than either bin-based method, particularly at small to intermediate bins (≤5 ms). Error bars indicate mean with 95% CI. Lines at the top indicate significance (*, p < 0.05). (I) A power analysis further revealed that *latenZy* required fewer neurons to detect latency differences between V1 and AM at α = 0.05 with 80% power, relative to bin-based methods using 5 ms bins. (J) This advantage held across a wide range of bin widths, highlighting *latenZy*’s increased sensitivity and robustness.

We applied *latenZy* to responses evoked by drifting gratings in 6,224 neurons (1,038 ± 237 neurons per area, mean ± SD) recorded across six visual cortical areas (Fig.5C, two example neurons). Consistent with prior findings (9,29) we observed a strong correlation between each area’s median response latency and its anatomical hierarchy score (r = 0.91, p = 0.013; Fig. 5D), validating *latenZy*’s ability to accurately recover meaningful inter-areal timing differences.

In agreement with earlier findings in both mice and macaques (8,9), response latencies within each visual area were broadly distributed (Fig. 5E). To compare latencies between areas, we computed latency dominance scores, which quantify how consistently neurons in one region respond earlier than those in another. For each area pair (reference and target), this score represents the difference between the proportions of neuron pairs where the reference neuron responded faster versus slower. Scores range from –1 (reference always slower) to +1 (always faster), with 0 indicating no consistent difference. For example, comparing LM (reference) to PM (target) yielded a dominance score of +0.24, indicating that LM neurons tended to respond earlier than those in PM (median 60.4 ms vs 68.2 ms, p < 2.1 × 10^−7^; Fig. 5E). We then examined a complete matrix of latency dominance cores across all area pairs (Fig. 5F) and correlated these with differences in anatomical hierarchy scores derived from connectivity patterns (Fig. 5B). This analysis revealed a strong relationship between functional latency and anatomical hierarchy (r = 0.79; Fig. 5G, left), which was notably stronger than correlations based on bin-based latency measures at an intermediate bin width (r = 0.50 for the half-max method and r = 0.64 for the SD-threshold methods; Fig 5G, middle and right). The findings suggest that dominance scores derived from *latenZy* estimates more effectively capture the temporal structure aligned with anatomical hierarchy than those based on bin-based methods.

To further assess the robustness of this result, we examined how varying bin widths impacted the relative performance of *latenZy* compared to bin-based methods. Using bootstrapping, we created distributions of mean correlation differences (*latenZy* minus each bin-based method) across a range of bin widths (Fig. 5H). *LatenZy* consistently outperformed the half-max method at all tested bin widths, with mean differences significantly greater than zero (p < 0.05). It also showed a significant advantage over the SD-threshold method at small to intermediate bin widths (≤ 5 ms), underscoring its improved temporal resolution.

Finally, we conducted a power analysis using Wilcoxon rank-sum tests to determine the number of neurons required to detect a significant difference in latency between V1 and AM, the areas with the greatest anatomical hierarchy separation (Fig. 5B). Tests based on *latenZy* required fewer neurons to reach 80% power at a 0.05 significance level compared to bin-based methods using 5 ms bins (Fig. 5I). This advantage of *latenZy* was apparent both at high temporal resolutions (≤5 ms) and at coarser time scales (>20 ms; Fig. 5J).

Together, these results demonstrate that *latenZy* not only captures biologically meaningful temporal structure in visual cortical responses but also offers clear advantages over traditional approaches—especially when high temporal precision or limited sample sizes are essential.

### *LatenZy2*: Estimating the onset of condition-specific spiking activity without binning

We introduced *latenZy* as a method to estimate the latencies of neural spiking responses to discrete events, like the onset of a visual stimulus. However, many studies require identifying when spiking activity begins to diverge between experimental conditions—for example, to pinpoint the onset of top-down modulation of activity in the visual cortex (39–41) or to determine the sequence in which choice-selective activity emerges across multiple brain regions (5,42). Standard approaches typically rely on binning spikes to create condition-specific PETHs, followed by statistical comparisons or decoding analyses. However, the reliance on binning and repeated testing can obscure fine temporal dynamics (20) and inflate false positives (43), making these methods poorly suited for precisely identifying when spiking rates begin to differ between conditions. To address this, we developed *latenZy2*, a method designed to estimate the latency of such spiking rate differences in a binning-free and statistically robust way. The latency estimate is refined iteratively, following a procedure similar to *latenZy* (Fig. 6A). A detailed description of the steps is provided in the Methods section, but we give a brief overview here.

**Figure 6.**
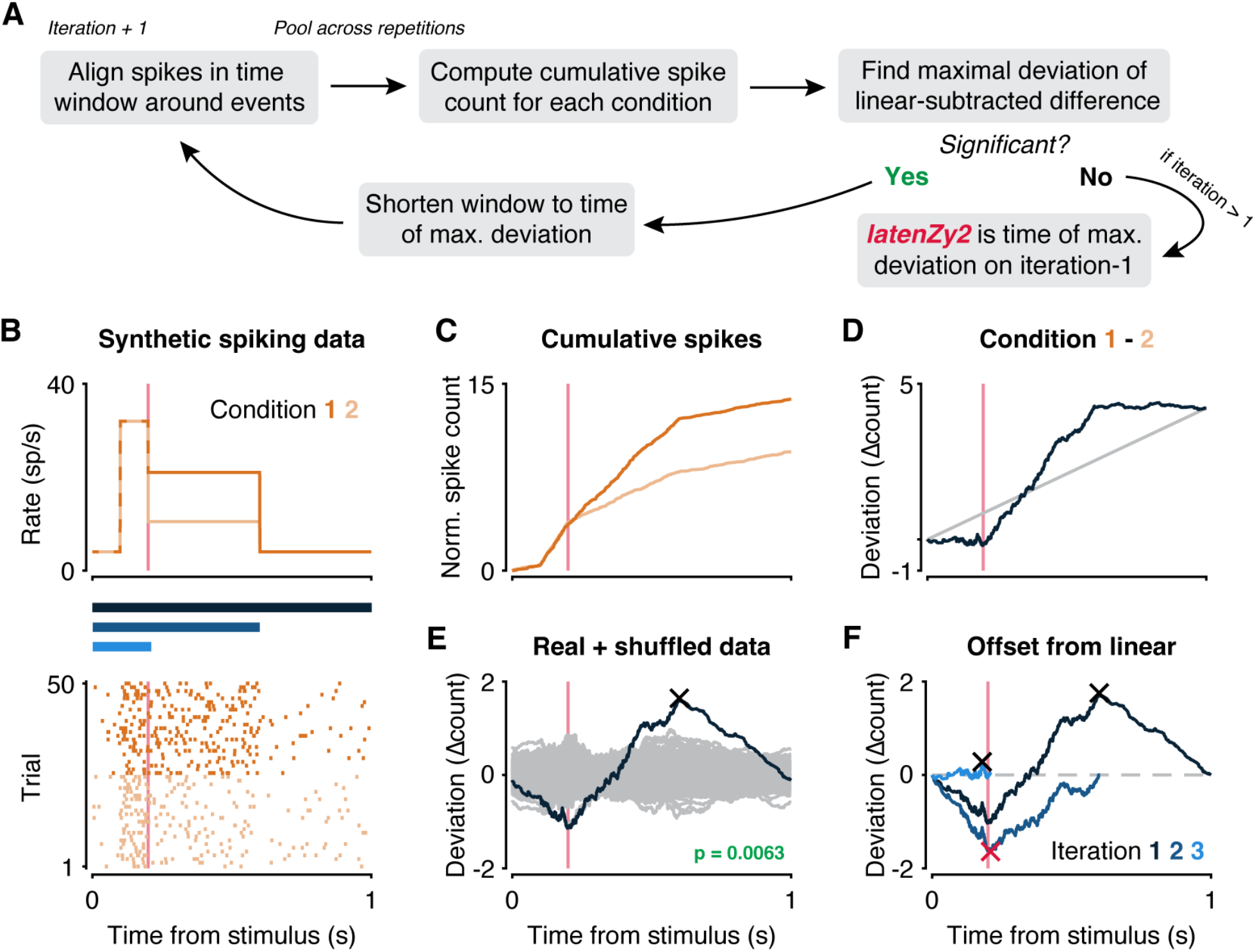
An overview of the *latenZy2* method. (A) A block diagram illustrating one iteration of *latenZy2*. (B) To illustrate its procedure, *latenZy2* was applied to spiking activity generated using an inhomogeneous Poisson process. The top panel shows the time-varying response functions used to drive the simulations: both conditions share an identical onset transient, but the sustained rate is higher for Condition 1 (dark orange) than for Condition 2 (light orange). The bottom panel shows raster plots of the resulting spike trains for each conditions, aligned to stimulus onset. Vertical pink lines indicate the point where the response functions start to diverge. Blue bars represent the time windows evaluated in each iteration of *latenZy2*. (C) Normalized cumulative spike counts are computed separately for each condition (dark and light orange curves). (D) Their difference yields a spiking-density deviation vector (dark blue curve), capturing the temporal evolution of the divergence between conditions. A linear baseline (gray line), representing a constant rate offset, is subtracted from the deviation vector to isolate the time-varying component. (E) The maximal deviation from the linear baseline (cross) is identified, and statistical significance is assessed by comparing to a null distribution generated from condition label-shuffled bootstrap iterations (gray curves). A p-value is computed based on the extremal deviations. (F) If the deviation is significant (p < 0.05), the analysis window is truncated at the latency corresponding to the maximal deviation, and the procedure is repeated. Iterations continue until no further significant divergence is detected. The red cross marks the final *latenZy2* estimate, indicating the earliest reliably detectable divergence in spiking between conditions.

We begin by aligning spikes within a window surrounding the event times for each condition (Fig. 6B) and pooling them across repetitions. Rather than comparing a neuron’s response to a baseline rate, we compute a temporal spiking-density deviation vector, defined as the difference between the normalized cumulative spike counts of the two conditions (Fig. 6C, D, dark blue curves). To isolate the time-varying component of this difference, we subtract a linear trend (Fig. 6D, gray line), which corresponds to a constant offset in spiking rates, and we identify the point of maximal deviation and its associated latency (Fig. 6E, cross). To assess the statistical significance of this deviation, we generate a null distribution using a repetition-swapping bootstrap procedure. In each bootstrap iteration, repetitions are randomly reassigned to one of the two conditions, a new deviation vector is computed (Fig. 6E, gray curves), and its extremal value is recorded. We then compute a p-value for the real data using a Gumbel distribution approximation (26). If the extremal deviation is statistically significant (p < 0.05, default cutoff), we narrow the upper bound of the analysis time window to the estimated latency and repeat the procedure (Fig. 6F). This process continues until no further significant divergence is detected. The final *latenZy2* estimate corresponds to the earliest time point— from the start of the analysis window—at which a significant difference in spiking rates between conditions can be reliably detected. This iterative, binning-free approach enables precise and statistically robust estimation of when neural responses begin to diverge between conditions. Next, we compare *latenZy2* to bin-based methods and showcase its use in examining attentional modulation in macaque visual cortex.

### *LatenZy2* outperforms binned methods in capturing top-down attention effects

Attentional modulation influences spiking activity throughout the visual hierarchy, but the causal flow of these changes remains debated. One leading hypothesis posits that selective attention operates via top-down feedback signals that increase the responsiveness of neurons processing attended stimuli. According to this view, the strongest and earliest attentional effects emerge in extrastriate visual areas, like V4, and then propagate backward to earlier areas such as V1 (44,45). Multiple studies support this idea, showing that attentional modulation progresses backward from extrastriate cortex to V1 (46), and that disrupting feedback from V4 eliminates attentional effects in V1 (47). Furthermore, attention-related spiking rate changes appear earlier in the superficial layers of V4 than in V1 (40), providing additional evidence for a top-down source.

To evaluate the utility of *latenZy2* in capturing these attentional dynamics, we applied it to a publicly available dataset (48) featuring simultaneous recordings from macaque V1 and V4 during a visual attention task. In this task, monkeys initiated a trial by fixating on a central spot while three colored square wave gratings appeared, one positioned within the receptive field of the recorded neurons (Fig. 7A). After a variable delay, a central color cue indicated the target grating to be attended. Subsequently, the monkeys had to detect and report a luminance change in the cued grating (not shown) to receive a reward, while ignoring dimming events in non-cued distractors. We applied *latenZy2* to estimate the onset of attention-related modulation by comparing spiking activity between two conditions: attend-toward (cue directs attention to the receptive field stimulus) and attend-away (cue directs attention elsewhere) (Fig. 7B).

**Figure 7.**
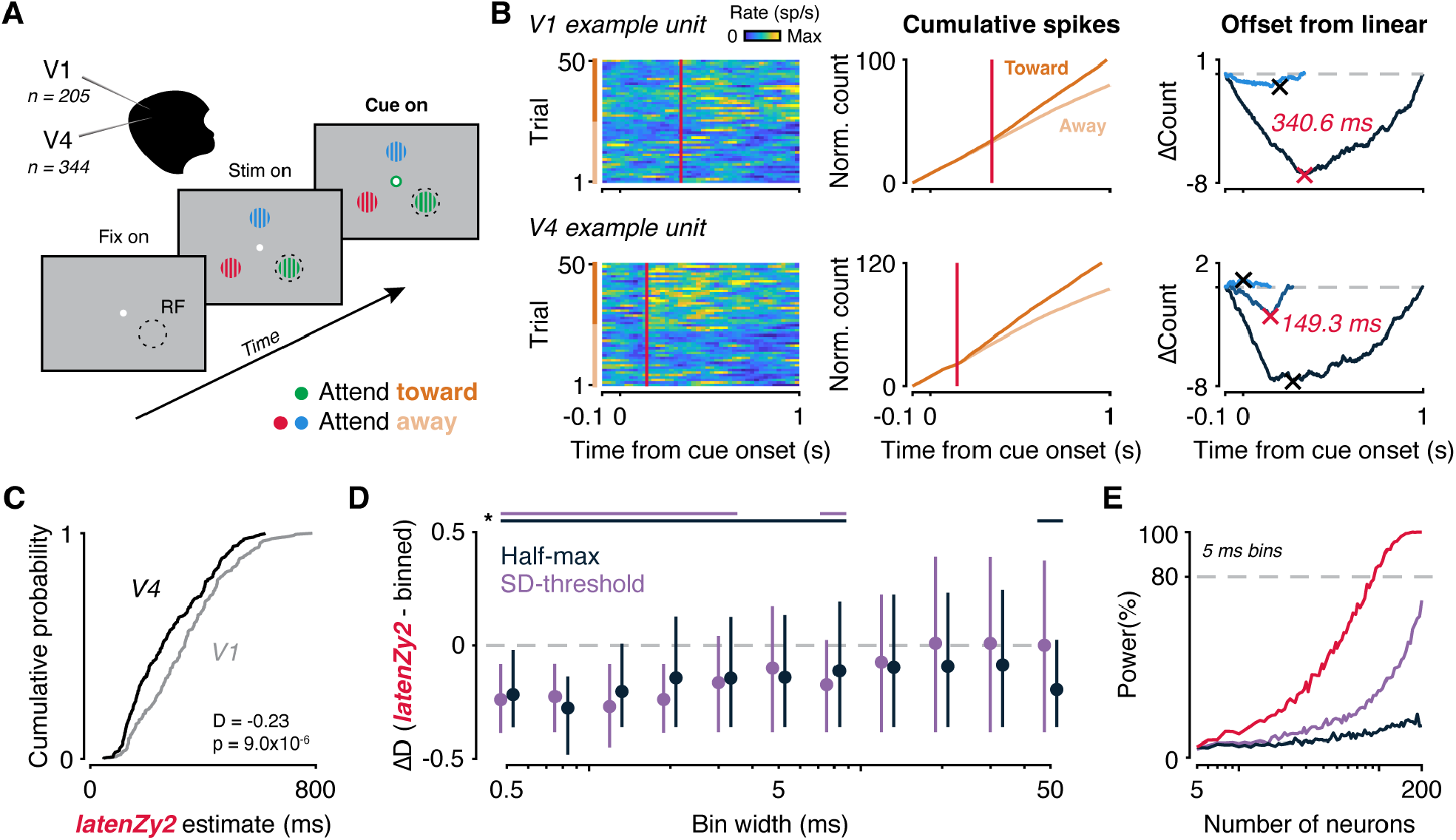
*LatenZy2* detects earlier attentional modulation in V4 compared to V1 in a visual attention task. (A) Illustration of the task design (48,57), in which monkeys fixated centrally while three colored gratings appeared, one positioned within the receptive field (RF) of the recorded neurons. After a delay, a central color cue indicated which grating should be attended. (B) Spike trains from example V1 (top) and V4 (bottom) units under attend-toward (cue to RF) and attend-away (cue elsewhere) conditions, aligned to cue onset (left). Trials are sorted by type: attend toward (dark orange) and attend away (light orange). Spikes were binned for visualization purposes only. *LatenZy2* was used to estimate the latency of attention-related modulation by identifying time-locked differences in spiking between conditions (middle, right). Red vertical lines and crosses indicate the *latenZy2* estimate for each unit. (C) At the population level, attentional effects were detected significantly earlier in area V4 than in V1 (median latencies: 255.2 ms vs. 334.7 ms; Wilcoxon rank-sum test, p = 9.0 × 10^−6^). A negative latency dominance score (–0.23) indicated that attentional modulation consistently emerged earlier in V4. (D) Compared to bin-based latency estimation methods (half-maximum and SD-threshold), *latenZy2* yielded more negative latency dominance scores at small to intermediate bin widths (0.5–8 ms). Lines at the top indicate significance (p < 0.05). (E) Power analysis showed that *latenZy2* achieved 80% power to detect the latency difference with a sample of 95 neurons per area at α = 0.05, whereas neither bin-based method reached this threshold at a 5 ms bin width within the range of sample sizes tested.

Consistent with prior work, *latenZy2* revealed significantly earlier attentional effects in V4 versus V1, with median latencies of 255.2 ms and 334.7 ms (Wilcoxon rank-sum test, p = 9.0 × 10^−6^; Fig. 7C). We further quantified the difference in the onset of attentional modulation between the V1 and V4 populations by computing a latency dominance score, scaled from −1 (all V4 latencies earlier) to +1 (all V1 earlier). The observed score of −0.23 indicates a consistent trend of earlier attentional modulation in V4 than in V1, supporting top-down feedback within the visual hierarchy.

To benchmark *latenZy2*, we compared it to two bin-based approaches—variations of the half-maximum and SD-threshold methods applied to the *difference* between the attend-toward and attend-away PETHs. Using a bootstrapping procedure, we generated distributions of latency dominance score differences between *latenZy2* and the bin-based methods (*latenZy2* minus bin-based) across multiple bin widths for statistical comparison. The advantages of *latenZy2* over bin-based methods were most pronounced at small to intermediate bin widths (0.5 to 8 ms), where it yielded significantly more negative latency dominance scores, particularly compared to the SD-threshold method (Fig. 7D). Finally, a power analysis using Wilcoxon rank-sum tests on latency estimates from V1 and V4 revealed that only tests based on *latenZy2* estimates reached 80% power at α = 0.05 with a sample of 95 neurons. In contrast, tests on bin-based estimates did not achieve this level of power at the tested sample sizes using a 5 ms bin width (Fig. 7E).

In conclusion, *latenZy2* is a binning-free, statistically robust method to precisely estimate when neural spiking rates diverge between conditions. It outperforms bin-based approaches in sensitivity and requires fewer neurons to achieve high statistical power. This makes *latenZy2* a powerful tool for studying the timing of neural response differences across various experimental contexts.

## Discussion

We introduced *latenZy*, a non-parametric, binning-free method for estimating neuronal response latencies, along with *latenZy2*, its extension designed to detect the onset of divergence in spiking activity between two conditions. These methods leverage cumulative deviations in spike timing to identify event-locked changes in trial-pooled neural activity, without requiring predefined time bins, arbitrary thresholds, or assumptions about response shape. This makes them particularly well-suited for analyzing neurons with diverse or idiosyncratic spiking patterns (Fig. 2), where standard methods often require *ad hoc* solution or optimization of parameters like bin widths, response threshold, or minimum response duration (17)—limiting reproducibility and comparability of latency estimates between neurons, areas, and studies.

We demonstrated that *latenZy* provides more precise and stable latency estimates than conventional bin-based techniques, especially at fine temporal resolution (Fig. 3). It reliably captured established physiological phenomena such as contrast-dependent latency shifts (Fig. 4) and hierarchical timing across mouse visual areas (Fig. 5). Power analyses using latency estimates derived from *latenZy* showed improved statistical power, requiring fewer neurons—and thus fewer animals—to detect significant latency differences in population analyses (Fig. 4, 5). *LatenZy2* further enables accurate estimation of when spiking activity begins to diverge between conditions, as shown in our analysis of attentional modulation in macaque V4 and V1. It revealed earlier divergence in V4, consistent with top-down feedback mechanisms, and outperformed bin-based methods in both sensitivity and sample size efficiency (Fig. 7).

Our work builds on earlier cumulative statistical methods for detecting neural response onsets. The cumulative sum method was originally applied to detect deviations in PETHs, using thresholds based on deviations from baseline spiking (49,50). These early approaches often lacked statistical grounding, relying on arbitrary thresholds to define onset. Later refinements introduced significance bounds based on point process theory, accounting for refractory effects in spike trains (51) but continued to rely on binned data and sensitive control parameters. Other advances improved temporal precision using second-order difference functions (52) or addressed non-uniform spiking expectations using Monte Carlo-based cumulative sum tests (53). More formal change-point detection approaches used maximum likelihood and least squares estimation under Poisson rate-change models (1). While these methods improved statistical rigor, they still required assumptions about rate structure and careful binning strategies. *LatenZy*, in contrast, preserves the temporal integration strength of cumulative approaches but eliminates the need for binning, stationarity, or rate modeling assumptions, enabling generalizable, high-resolution estimation across datasets.

Nonetheless, *latenZy* has limitations. It produces a single latency estimate per neuron by aggregating spikes across trials and therefore does not capture trial-to-trial variability in response timing. When within-neuron latency fluctuations are of interest, alternative methods may be more appropriate (13,17,54,55). Additionally, the statistical test underlying *latenZy* may fail to reach significance when responses are weak, variable, or inconsistent, potentially excluding low-firing or subtly modulated units. However, this trade-off reduces false positives and enhances robustness at the population level, supporting reliable population-level inference. Importantly, prior studies have benchmarked the statistical framework that *latenZy* employs (26,27), demonstrating that it includes more neurons than alternative methods at a similar false positive rate. A limitation of *latenZy2* is that it is designed for pairwise comparisons. While sufficient for many experimental designs, it does not extend directly to comparisons across more than two conditions. In such cases, multivariate techniques or decoding-based frameworks may be more suitable for characterizing the full structure of condition-dependent temporal dynamics.

Overall, *latenZy* offers a compelling combination of statistical rigor and practical usability. It operates directly on spike times without requiring user-defined parameters and is broadly applicable to both single-condition and two-sample latency estimation tasks. Together, *latenZy* and *latenZy2* provide robust, binning-free, and biologically grounded tools for precise temporal analysis of neural spiking, especially in high-throughput experimental settings. Open-source implementations are available in both Python and MATLAB at: https://github.com/Herseninstituut/latenzy.

## Methods

### LatenZy

*LatenZy* is an extension of the Zenith of Event-based Time-locked Anomalies (ZETA) test (26,27) and is designed to estimate neural response latencies. Building on the ZETA framework, we compute the *latenZy* estimate, denoted as *τ*_*Z*_, on a vector of *i =* [1 … *n*] spike times ***x***, and a vector of *k =* [1 … *q*] event times ***w***. We perform this calculation within a time window [*t*_0_, *t*_1_] around each event time, with boundaries initially set to satisfy either *t*_0_ ≤ 0 and *t*_1_ > 0, or *t*_0_ < 0 and *t*_1_ ≥ 0. To ensure accurate estimation, the window [*t*_0_, *t*_1_] must exclude confounding time-locked activity, such as offset responses to preceding stimuli.

We define the response latency as the earliest time after *t*_0_ at which a significant time-locked deviation in repetition-pooled spiking activity is first observed. This time point is identified using an iterative procedure that progressively refines the estimate across successive steps.

### Stitching data across event windows

In each iteration, we start with a “data-stitching” step, where we remove any data points that fall outside the event-window bounds for each repetition (27). This guarantees that only spikes within the relevant time windows are included, maintaining consistency between the real data and the randomly resampled data used for the statistical comparison described later. Specifically, we remove data points *x*_*i*_ between event *k* and *k* + 1 for which:

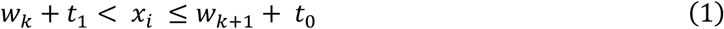

To account for the excluded periods, we adjust both the spike times and event times. The adjusted spike times *x*_*i*_^*\**^ and event times *w*_*k*_^*\**^ are computed as:

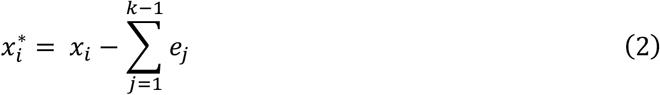

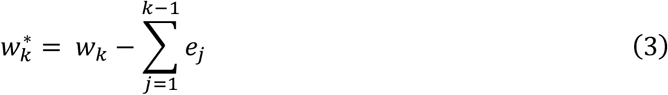

Here, *e*_*j*_ represents the duration of the excluded interval between repetitions, defined as:

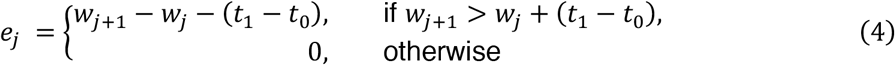

For notational simplicity, from this point onward, we refer to the adjusted spike and event times, ***x***^*\**^ and ***w***^*\**^, simply as ***x*** and ***w***.

### Computing the maximal deviation from uniform spiking

After the data-stitching step, we align the spike times to the event times by computing the relative spike times within the defined time window [*t*_0_, *t*_1_] around each event. For each spike *x*_*i*_ that falls within this window relative to event *w*_*k*_, we calculate the relative spike time ***v***_***i***_ as:

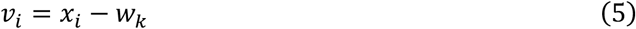

This produces a single vector ***v*** of aligned spike times aligned to the events. To ensure that the full epoch is covered, we add two artificial spikes at *t*_0_ and *t*_1_. Next, we sort the spike times in ***v*** such that *v*_*i*_ *< v*_*i*+1_ and compute the fractional position *g*_*i*_ of each spike time in ***v***:

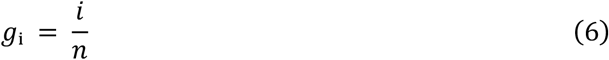

where *n* is the total number of spike times in ***v***. The vector ***g*** represents the cumulative density function of spikes sampled at the spike times in ***v***.

To assess whether the spiking activity is time-locked to the event, we compare this to a linear baseline density vector ***b***. Under the null hypothesis of a constant spiking rate (i.e., no modulation by the event) the cumulative density function should be linear. Therefore, as the number of events *q* increases, the expected fractional position of spike *i* at time *v*_*i*_ converges to the spike time divided by the window duration:

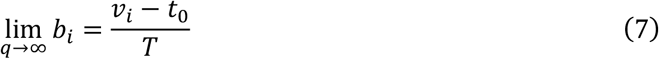

where *T* = *t*_1_ – *t*_0_ is the total duration of the window.

The difference *δ*_*i*_ between *g*_*i*_ and *b*_*i*_ thus gives a neuron’s deviation from a temporally non-modulated spiking rate at each time point *v*_*i*_:

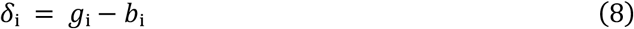

Sudden changes in the slope of the deviation vector ***δ*** reflect changes in the trial-pooled spiking rate. In each iteration, we extract the extremal value of ***δ***, denoted *ζ*_*r*,_ which corresponds to the point of maximum absolute deviation from the expected uniform spike density:

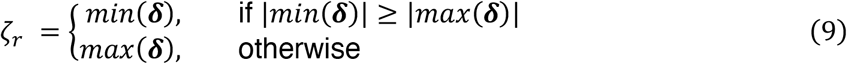

The associated latency *τ*_*ζ*_ is determined from ***v***:

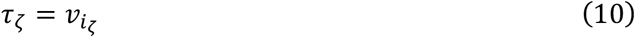

where *i*_*ζ*_ is the index of the spike time associated with the maximal absolute deviation *ζ*_*r*_.

### Testing the maximal deviation against a null distribution

Having computed the maximal deviation *ζ*_*r*_, we seek to quantify its statistical significance. To do so, we construct a null hypothesis distribution by repeating the above procedure *M* times with jittered event times *w’*. For each jitter iteration *m* and event *k*, we randomly shift each event time by a random amount sampled from the interval [-*T, T*]:

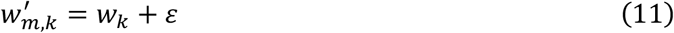

Where each *ε* is independently drawn from a uniform distribution *U*:

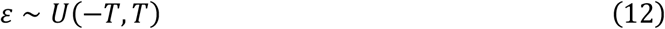

For each jitter iteration, we compute the deviation vector ***δ****’(m)*:

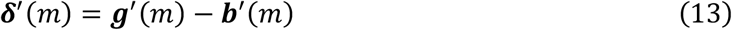

From this, we extract the extremal value *ζ’(m)*, analogous to *ζ*_*r*_:

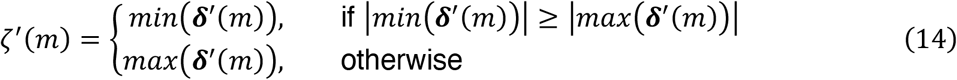

Next, we create a mean-normalized version of *ζ*_*r*_:

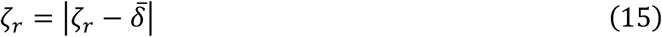

where the mean deviation is computed as:

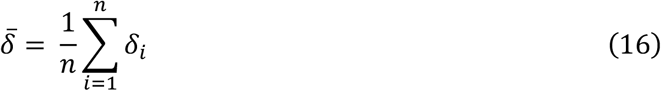

Similarly, for each jitter iteration *m*, we compute the mean-normalized extremal deviation:

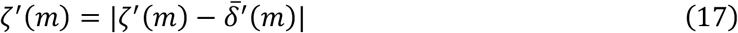

We can now compute the p-value of *ζ*_*r*_ by determining its quantile position in the set of *M* samples *ζ’*, or by approximating the distribution of *ζ’* with the Gumbel distribution (26). Both approaches provide an estimate that converges asymptotically as the number of jitter iterations *M* increases.

To obtain a corrected metric *ζ* that is interpretable as a z-score, we apply the inverse cumulative normal distribution *φ*^−1^ to the tail probability of *ζ*_*r*_:

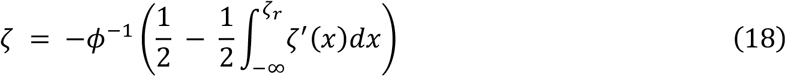

### Iterative refinement of the response latency estimate

In each iteration *l*, we evaluate the p-value of the extremal deviation *ζ*_*l*_ against a predefined threshold (*α* = 0.05, default). If the p-value is below this threshold indicating statistically significant time-locked spiking, the analysis window is narrowed to [*t*_0_, *τ*_*ζl*_] and the next iteration *l* + 1 is performed by repeating the steps above.

If, instead, the p-value is equal to or exceeds the cutoff threshold, suggesting that no further significant modulation is present within the window, the response latency *τ*_*Z*_ is defined as the value obtained in the previous iteration (*l* – 1):

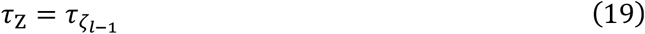

When this stopping condition arises during the first iteration (i.e., *l* = 1), it is unlikely that the neuron’s spiking activity exhibits any significant event-related modulation within the examined time window, and *τ*_*Z*_ remains undefined.

### LatenZy2

In many experimental paradigms, a key question is *when* spiking responses begin to diverge between two conditions. To address this, we developed *latenZy2*, a two-sample extension of *latenZy*, designed to estimate the earliest time point at which spiking rates differ significantly. The only requirement we place on the data is that the temporal window [*t*_0_, *t*_1_] is identical for both conditions, with the same boundary restrictions as *latenZy*.

### Computing the maximal difference in spiking between conditions

The *latenZy2* estimate, denoted as *τ*_*Zd*_, is computed on the event-relative spike times ***v***^*α*^ of condition *α* and ***v***^*β*^ of condition *β*. These event-relative spike times are obtained by using Equation 5 on a vector of spike times ***x*** and event times ***w***^*α*^ and ***w***^*β*^ for each condition and sorted so that *v*_*i*_ < *v*_*i*+1_. Note that the *α* and *β* superscripts are condition indicators, not exponents.

Collapsing all the spikes across trials without normalization would lead to differences in the total spiking numbers when there is a difference in the number of trials per condition but not spiking rate. To avoid this, we construct a cumulative spiking vector for each condition based on the average number of spikes per event window:

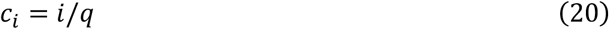

Where *q* is the number of events. We compute this separately for both conditions, obtaining ***c***^*α*^ and ***c***^*β*^. To compare these vectors, we first create a reference time vector ***ρ*** containing all the spike times of ***v***^*α*^ and ***v***^*β*^:

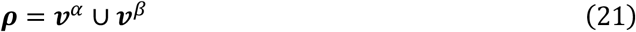

We then linearly interpolate the cumulative spike vector ***c***^*α*^ onto the time points in ***ρ***. For each time point *ρ*_*i*_, the interpolated value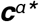:

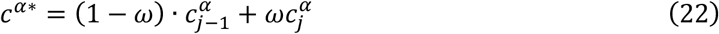

Where *ω* is the interpolation weight defined by:

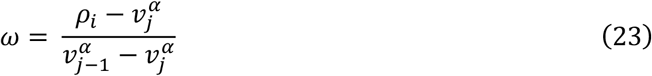

We apply the same procedure to 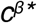 for condition *β*. Now, we compute the difference in cumulative spike counts between the two conditions as:

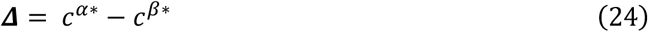

To remove any linear offset reflecting a constant offset in spiking rates rather than transient divergence, we detrend ***Δ*** by subtracting a linear baseline connecting its endpoints:

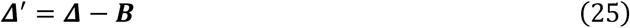

Where the linear baseline ***B*** is computed as:

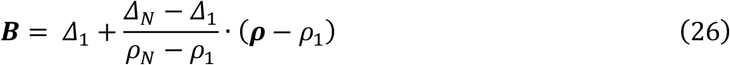

Here, *ρ*_*1*_ and *ρ*_*N*_ denote the first and last time points in the combined spike time vector ***ρ***, and *Δ*_*1*_, *Δ*_*N*_ are the corresponding values of ***Δ***.

Next, we extract the extremal value of ***Δ***’, denoted as *ζ*_*dr*_, following the procedure in Equation 9. The corresponding latency is given by:

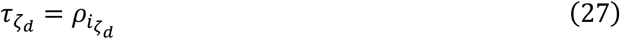

where *i*_*ζd*_ is the index of the spike time associated with the maximal absolute difference *ζ*_*dr*_.

### Testing the maximal difference against a null distribution

To construct a null hypothesis distribution for *latenZy2*, we begin by pooling all trials from conditions *α* and *β* into a unified set. For each event *k*, we collect the spikes occurring within the window [*t*_0_, *t*_1_] relative to that event by defining the set:

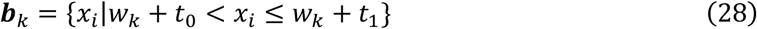

We then align these spikes to the event time by subtracting *w*_*k*_:

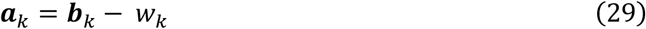

The union of these aligned spike sets across all events in both conditions forms the pooled set:

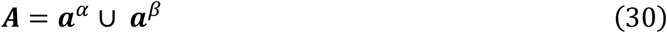

To create a random null-hypothesis sample *m* for condition *α*, we select *q*^*α*^ random sets from ***A*** without replacement and concatenate them:

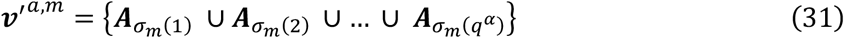

Here, ***σ***_*m*_ is a vector containing the *q*^*α*^ random integers in the range [1, *q*^*α*^ *+ q*^*β*^]. We repeat this procedure to generate shuffle-randomized spike vectors ***v****’*^*β,m*^ for condition *β*.

Each pair (***v****’*^*α,m*^, ***v****’*^*β,m*^) is then used to compute a null-hypothesis sample maximal deviation *ζ*_*d*_’(*m*) by applying Equations 20-26 and 14 to the shuffled data. By applying the normalization and transformation steps analogous to those used in *latenZy* (Equations 15-18), the null samples *ζ*_*d*_’(*m*) are converted into a statistical metric and p-value.

Finally, as with *latenZy*, we iteratively refine the latency estimate *τ*_*Zd*_ by narrowing the analysis window until the extremal deviation fails to reach statistical significance.

### Bin-based latency estimation methods

We benchmarked *latenZy* against two widely used bin-based latency estimation methods: the half-max method (15,16) and the standard deviation (SD)-threshold method (13). Both methods operate on event time histograms (PETHs), which we computed by binning spiking activity. We used 11 logarithmically spaced bin widths ranging from 0.5 ms to 50 ms. To enable comparison with *latenZy2*, the bin-based methods were applied to the difference PETH—that is, the binwise difference between the condition-specific PETHs. Baseline activity was defined as all activity occurring before stimulus onset (*t* < 0), but within the defined analysis window.

#### Half-max method

We defined the latency as the center of first bin after stimulus onset in which the PETH deviates from baseline by at least half the peak response magnitude (positive or negative). Let *λ(t)* denote the PETH at time *t* and let *λ*_*B*_ be the mean of the PETH during the baseline period. The peak response is defined as:

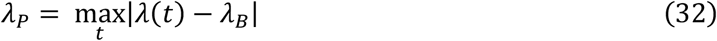

The latency then corresponds to the first bin after stimulus onset satisfying:

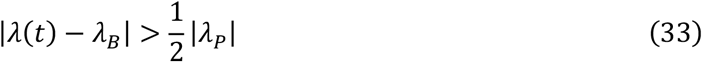

#### SD-threshold method

We defined the latency as the first bin after stimulus onset in which the PETH exceeds the baseline mean, *λ*_*B*_, by more than *k* standard deviations:

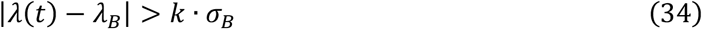

Where *σ*_*B*_ is the SD across baseline time bins of the trial-averaged PETH—reflecting temporal variability in the response, not trial-to-trial variability. We used *k* = 2.58, corresponding to a two-tailed significance level of 0.01 (32,33).

### Experimental data for benchmarking

The electrophysiological data sets used in this study have been previously described elsewhere (9,26,48,56,57). Here, we summarize the aspects relevant to the current analyses. In all cases, units with very low spiking activity (<0.1 sp/s) or high non-stationarity were excluded. For benchmarking, *latenZy* estimates were constrained to be positive, as both the half-max and SD-threshold are by definition limited to post-stimulus responses. This constraint was appropriate, as we did not expect meaningful latencies to occur before stimulus onset.

#### Zebrafish retina recordings (26)

Extracellular recordings of retinal ganglion cell activity were obtained from isolated retinas of adult zebrafish (*Danio rerio*) using a 60-electrode array. Neural signals were recorded at 25 kHz, bandpass filtered, and sorted offline into single-unit activity based on spike waveform features. Visual stimulation consisted of a 0.5 s full-field white light flash, preceded by 0.5 s and followed by 1.0 s of darkness. The stimulus was projected onto the photoreceptor layer and repeated 50 times. The analysis window for *latenZy* was set from −0.1 to 0.5 s relative to flash onset.

#### V1 Neuropixels recordings (56)

Neuropixels recordings were performed in awake C57BL/6J mice, targeting various brain regions across 21 sessions. Spike sorting was performed using Kilosort, and probe locations were registered to the Allen Common Coordinate Framework. All 16 recordings that included penetrations of the primary visual cortex (V1) were included in the analysis. Visual stimuli consisted of full-field drifting gratings presented in 24 directions (15° increments), each repeated 20 times. Spiking activity was pooled across directions and analyzed within a time window from −0.1 to 1.0 s relative to stimulus onset.

#### Allen Brain Institute dataset (9)

The Allen Brain Institute’s Visual Coding Neuropixels dataset (Functional Connectivity subset; Siegle et al., 2021) includes simultaneous single-unit recordings from 4 to 6 visual cortical areas—V1, RL, LM, AL, PM, and AM—in 26 awake mice. Visual stimuli were drifting gratings moving in four directions (−45°, 0°, 45°, and 90°), with 0° corresponding to rightward motion. For the contrast sensitivity benchmark, V1 responses to gratings at five contrast levels (0.13, 0.2, 0.35, 0.6, and 1.0) were analyzed within a −0.1 to 0.5 s window relative to stimulus onset, with 60 repetitions per contrast level (15 per direction). Power analysis compared responses to low (0.1) and high (0.8) contrast stimuli over a longer −0.2 to 2.0 s window, using 300 repetitions per contrast. The hierarchy benchmark assessed responses to 0.8 contrast gratings across all six visual areas within the same −0.2 to 2.0 s window, using 300 total trials (75 per direction). See https://portal.brain-map.org/circuits-behavior/visual-coding-neuropixels for more information.

#### Macaque V1/V4 recordings during attention task (48,57)

Data were reanalyzed from three adult male rhesus macaques (*Macaca mulatta*) trained on a selective visual attention task. Each animal was implanted with a head post and recording chamber over visual areas V1 and V4. Visual stimuli were presented on a CRT monitor, and receptive fields were mapped using reverse correlation with flashed black squares. During the task, monkeys fixated while three colored gratings were presented and were rewarded for responding to dimming of the target stimulus while ignoring distractor dimming. Neural recordings were made using 16-contact laminar silicon probes across 83 sessions, with signals sampled at 32.7 kHz. Spikes were extracted via common average referencing and thresholding to obtain multiunit activity (~100 Hz per channel during task epochs). Analyses focused on the −0.1 to 1.0 s window around cue onset.

## Statistics

### Monotonicity score

To quantify the directional consistency of latency changes across stimulus contrasts for a single neuron, we computed a monotonicity score. Given a latency vector ***A*** = {*a*_1_, *a*_2,_ *…, a*_*N*_} ordered by increasing contrast, the score *M* is defined as:

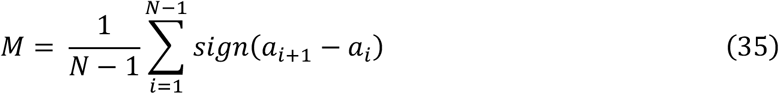

where *sign(x)* ∈{−*1, 0, +1*} denotes the sign of the difference: +1 for increases, −1 for decreases, and 0 for no change. The score ranges from −1 (strictly decreasing) to +1 (strictly increasing), with 0 indicating no consistent trend.

#### Latency dominance score

To assess whether one brain area consistently exhibits shorter or longer latencies compared to another, we compute a latency dominance score. For two sets of latency measurements, ***A*** = {*a*_1_, … *a*_*n*_} and ***B*** = {*b*_1_, … *b*_*m*_}, the score *D* quantifies the balance between how often latencies in ***A*** exceed those in ***B*** versus the reverse:

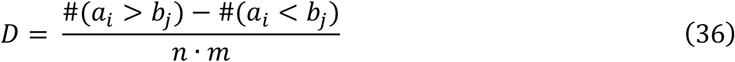

Where #(*a*_*i*_ *> b*_*j*_) is the count of all pairwise comparisons in which values from ***A*** are greater than those from ***B***, and similarly for #(*a*_*i*_ *< b*_*j*_). The score ranges from −1 (all latencies in ***A*** are shorter than in ***B***) to +1 (all latencies in ***A*** are longer), with zero indicating no consistent latency difference.

#### Estimating variability of latency estimates

To assess the reliability of *latenZy* and other latency estimation approaches, we implemented a subsampling procedure where we randomly selected half of the available trials (240 from 480 total) across 250 iterations, then computed the standard deviation of latency estimates across these iterations. We chose subsampling without replacement over conventional bootstrapping to avoid creating superimposed spikes that could introduce artifactual peaks in *latenZy*’s cumulative deviation calculations. Our analysis included only units that yielded valid *latenZy* estimates in at least 70% of subsamples to ensure result stability.

For the bin-based methods, we added half the bin width to the standard deviation of the latency estimates to reflect the inherent temporal uncertainty introduced by binning. Since binning limits the temporal resolution of latency estimates, this adjustment accounts for the fact that the true latency could fall anywhere within the selected bin.

#### Bootstrapping approaches

We employed non-parametric bootstrapping to evaluate differences between *latenZy* and bin-based latency methods across multiple performance metrics: latency estimates, estimation variability, monotonicity scores, and dominance scores. For each metric, we calculated values for all recorded neurons using both approaches and computed paired differences (*latenZy* values minus bin-based values). We then created bootstrap distributions of mean difference by resampling these *N* paired differences with replacement across 10,000 iterations (where *N* represents the total number of neurons/units). Statistical significance was assessed using two-tailed tests for latency estimate comparisons and one-tailed tests for all other metrics, with no correction for multiple comparisons.

For dominance score comparisons, we conducted bootstrapping at the brain area level using two complementary approaches. In the first analysis, we resampled latency values within each brain region, recalculated dominance scores for each bootstrap iteration, and examined correlations with established anatomical hierarchy rankings. Statistical significance was determined by bootstrapping the paired differences in Pearson correlation coefficients. In the second analysis, we directly contrasted dominance scores computed on *latenZy2* estimates versus bin-based estimates, using bootstrap resampling of paired differences to test whether *latenZy2* consistently produced more extreme dominance scores compared to alternative approaches.

#### Power analyses

To evaluate and compare the sensitivity of multiple latency estimation methods, we conducted power analyses based on pairwise statistical comparisons. For each method and bin width, latencies were estimated across either different contrast levels or cortical areas. We applied Wilcoxon signed-rank tests for within-subject (contrast) comparisons and Wilcoxon rank-sum tests for between-area comparisons. To assess each method’s ability to reliably detect latency differences, we performed 1,000 Monte Carlo simulations per sample size for each comparison. In each simulation, samples of a given size were randomly drawn from the empirical latency distributions, and the corresponding statistical test was applied at a significance level of α = 0.05. Statistical power was defined as the proportion of simulations in which the test yielded a significant result. We considered a method to demonstrate sufficient sensitivity if power exceeded 80%. This procedure allowed us to determine the minimum sample size at which each latency estimation method achieved reliable statistical detection of latency differences.

## Data availability

Source data for this study are openly available at https://portal.brain-map.org/circuits-behavior/visual-coding-neuropixels (Allen Brain Institute Visual Coding Neuropixels dataset) https://doi.gin.g-node.org/10.12751/g-node.b0mnn2/ (macaque V1/V4 recordings). Other data will be made available upon reasonable request.

## Code availability

Open-source code for the MATLAB and Python implementations of *latenZy* and *latenZy2* can be found here: https://github.com/Herseninstituut/latenzy.

## Acknowledgements

R.H. was supported by the NWO Open Competitie ENW grant M20.114. We would like to thank Eric Lowet for testing an early version of the code and Jorrit Montijn for useful discussions.

